# Improving bacterial metagenomic research through long read sequencing

**DOI:** 10.1101/2023.10.31.564966

**Authors:** Noah Greenman, Sayf Al-Deen Hassouneh, Latifa S. Abdelli, Catherine Johnston, Taj Azarian

## Abstract

Metagenomic sequencing analysis is central to investigating microbial communities in clinical and environmental studies. Short read sequencing remains the primary data type for metagenomic research, however, long read sequencing promises advantages of improved metagenomic assembly and resolved taxonomic identification. To assess the comparative performance of short and long read sequencing data for metagenomic analysis, we simulated short and long read datasets using increasingly complex metagenomes comprised of 10, 20, and 50 microbial taxa. In addition, an empirical dataset of paired short and long read data from mouse fecal pellets was generated to assess feasibility. We compared metagenomic assembly quality, taxonomic classification capabilities, and metagenome-assembled genome recovery rates for both simulated and real metagenomic sequence data. We show that long read sequencing data significantly improves taxonomic classification capabilities and assembly quality. For simulated long read datasets, metagenomic assemblies were completer and more contiguous with higher rates of metagenome-assembled genome recovery. This resulted in more precise taxonomic classifications. Analysis of empirical data demonstrated that sequencing technology directly affects compositional results. Overall, we highlight strengths of long read sequencing for metagenomic studies of microbial communities over traditional short read approaches. Long read sequencing improved the accuracy of classification and abundance estimation. These results will aid researchers when considering which sequencing platforms to use for metagenomic projects.

**Data description:** The experimental metagenomic sequence data used for comparison of short and long read data from the same source are available from NCBI’s Sequence Read Archive (SRA) under Bioproject accession ID PRJNA1092431.

## Introduction

Metagenomics has enabled the study of microbial communities for agricultural, ecological, and clinical applications without the need for isolation or lab cultivation of bacteria (Chen and Pachter 2005; Cheng et al. 2019; Song et al. 2022; Suttner et al. 2020). Advancement in the field has been driven by the continued improvement of sequencing technologies, which have enabled generation of high-quality and low-cost genomic data for studying microbial populations with greater resolution (Wallen et al. 2022). Currently, the most widely used next-generation sequencing (NGS) technology are short-read sequencers, such as those from Illumina™. Short-read sequencing platforms generate 75-300 bp single or paired-end reads with per-base accuracy estimated at 99.9% (Latorre-Pérez et al. 2020). Before the recent reduction in cost attributed to the widespread availability of NGS technology, so-called microbiome studies often employed metabarcoding, where small portions of the 16S rRNA gene is sequenced for taxonomic classification (Begmatov et al. 2022; Oyserman et al. 2022; Sun et al. 2022). Although effective for genus level classification, metabarcoding provides limited species level resolution (Gupta et al. 2019). Where metabarcoding uses small regions of a single gene, shotgun metagenomics involves the sequencing of whole genome fragments, which can be used to reconstruct partial or whole microbial genomes through reference-based or *de novo* assembly approaches. Offering greater taxonomic resolution, *de novo* assembly is particularly useful for the identification of species that are non-culturable or lack robust representation in genomic databases. Shotgun metagenomic sequencing can also characterize variation in gene content, single nucleotide polymorphisms, and mobile genetic elements (Berglund et al. 2019; Sanderson et al. 2020).

In recent years, 3^rd^ generation “long-read” sequencing platforms such as those from Oxford Nanopore Technologies™ (ONT) and PacBio have increasingly been applied to clinical and biomedical research (Charalampous et al. 2019; Sanderson et al. 2020). These platforms use a single-molecule sequencing approach that generates significantly longer DNA sequences (Petersen et al. 2019). A major strength of 3^rd^ generation sequencers is their ability to span regions of genomes that are difficult for short-read sequencers to resolve, including common repetitive elements and other low complexity regions (Koren and Phillippy 2015). Functional analyses are further improved through the use of long reads as they can capture whole genes and preserve genetic structures (Singleton et al. 2021). Although there are numerous benefits, some notable drawbacks of long-read sequencing include lower per base accuracy. However, recent versions promise improvement of accuracy to 99% (Amarasinghe et al. 2020; Sereika et al. 2022). In addition, available tools for long-read data analysis are relatively limited when compared to tools for analysis of short-read data. More recently, the availability of tools has improved, with the number long-read sequencing programs more than quadrupling since 2018 (Amarasinghe et al., 2021, 2023).

Due to its high accuracy, wide-spread availability, and well-established published pipelines, short-read sequencing is typically employed over long read sequencing for metagenomic studies (Govender et al. 2021). However, the advantages afforded by long-read sequencing makes it an approach particularly well-suited for metagenomic research. In this study, we sought to investigate the relative performance of short- and long-read sequencing for metagenomic analysis. To this end, we simulated metagenomic data from synthetic microbial communities of increasing complexity and measured precision, recall, and F-score. We then applied paired short- and long-read sequencing in parallel to mouse fecal samples, and empirically compared the relative performance.

## Materials and Methods

### Simulated metagenomes of synthetic microbial communities

Nine data sets of simulated metagenomic sequencing reads were generated (Figure 1). Each set consisted of 10 unique, simulated metagenomic samples of varying complexity. Complexity was defined by the number of comprising taxa, the relative abundance of organisms in a metagenome, and the presence or absence of simulated sequencing errors. The size of a simulated metagenome consisted of genomes from 10, 20, or 50 taxa with abundances either evenly distributed or randomly generated and simulated sequencing errors present or absent. The three types of sequence data included “Perfect”, “Uneven”, and “True” datasets. Perfect datasets consisted of reads with no errors and even organism abundance distribution, uneven datasets without errors and random abundance values for each organism, and true datasets with simulated errors specific to each sequencing platform for a given sequence type and random abundance values for each organism. Simulated long-reads were generated to reflect sequence data from ONT’s MinION™ and short-reads mirrored sequence data from Illumina’s™ HiSeq™ sequencer.

**Figure 1.**
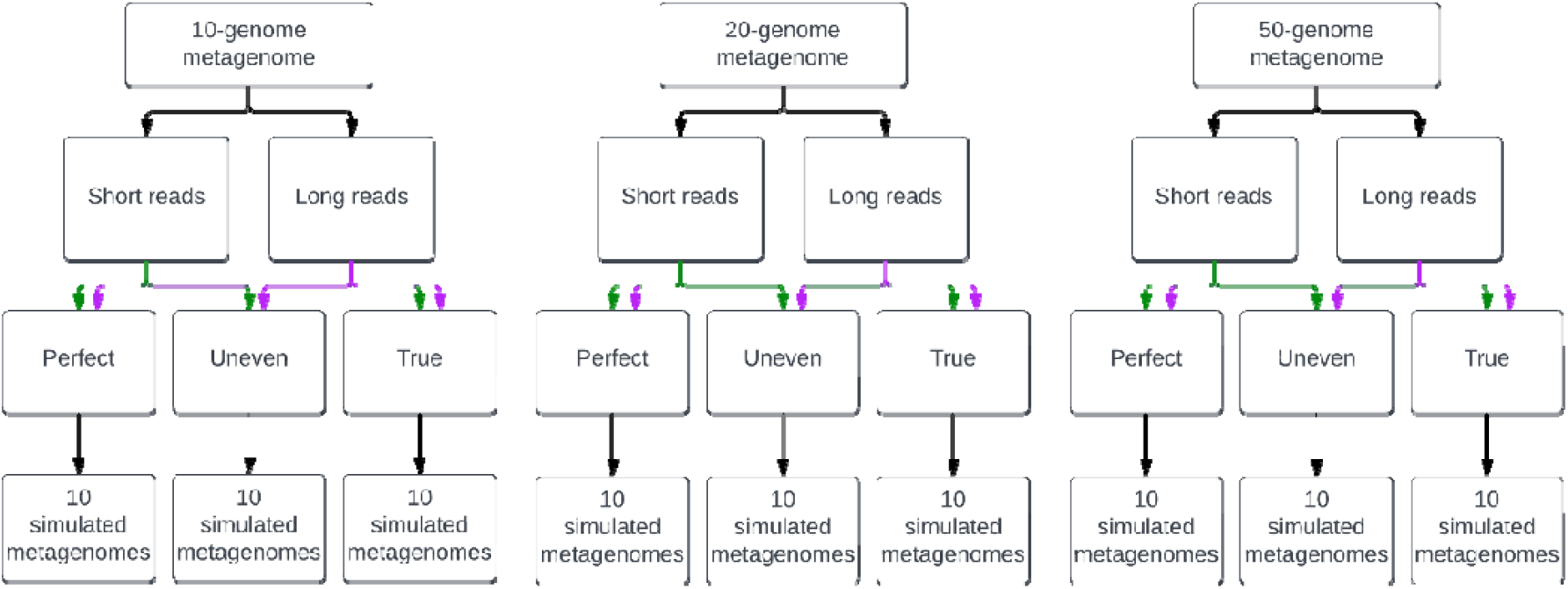
Simulation of metagenomic datasets of increasing complexity. “Perfect” metagenomic samples were simulated with no sequencing errors and an even abundance distribution of organisms. “Uneven” samples with no sequencing errors and randomly assigned abundance values for organisms in the metagenome, and “True” datasets with sequencing errors specific to each read type based on the platform they would be generated from and randomly assigned abundance values for members of each metagenome.

Construction of synthetic metagenomes was performed using an assembly structure report downloaded from NCBI (Downloaded 04/18/2022; Figure S1). This report contained all bacteria present in RefSeq with the following criteria for inclusion: 1) Genomes must be part of the latest RefSeq database, 2) they must have a complete genome assembly, and 3) all anomalous assemblies were excluded.

Simulation of ONT sequence data was performed using NanoSim version 3.0.0 (Yang et al. 2021). NanoSim input files (genome, abundance, and DNA lists) were created using a custom python script specifying parameters according to the level of metagenome complexity. The genome list (-gl) contained randomly selected taxa from the assembly structure report. The number of taxa selected was based on the desired size of the metagenome: 10, 20, or 50. Similar organisms could be present in a metagenome, such as different strains of the same species. The abundance list (-al) contained each organism and a value defining the abundance of that organism in the metagenome. This value was either equal across all organisms or randomly generated such that the total abundance of all organisms was 100%. The DNA list (-dl) contained all genetic molecules present in each organism’s genome. This included an organism’s chromosome and any plasmids, as well as whether the molecule was circular or linear. For simulating errors (-c), the pre-trained model ‘metagenome_ERR3152364_Even’ supplied by NanoSim was used. The basecaller (-b) specified was *guppy*. The ‘--perfect’ option was specified for datasets without simulated errors, including “Perfect” and “Uneven” datasets. This option was not used for “True” datasets. All simulated reads were generated in FASTQ format (--fastq). The total number of reads generated per metagenome was 650,000, with reads having an N50 of approximately 5.2 Kbps. Total bases for each sample were approximately 2.67 Gbps. These values were selected to mimic realistic outputs from nanopore sequencing. For long reads, filtering was performed using Filtlong version 0.2.1 (R. Wick 2021). Long reads were filtered by minimum read length of 1000 bp (--min_length 1000).

Simulation of short read sequence data from an Illumina™ HighSeq™ was done using the randomreads.sh script from BBMap version 37.93 (Bushnell 2020). Ten million paired-end reads were generated for each simulated metagenome. When creating metagenomic short read data, the abundance of each genome was determined from the abundance list used for simulating long reads. Dividing 10 million by the abundance for each genome yielded the number of short reads to simulate. Each iteration of randomreads.sh took a single genome and the number of reads specific to that genome and output a FASTQ file containing interleaved paired-end short reads. All datasets consisted of paired-end reads 150 bp in length (paired=t, len=150), named according to their genomic origin and without coordinates (simplenames=t), read quality was set to 36 (q=36), and a single insert size distribution was specified (superflat=t). Error rates were controlled by specifying the maximum number of SNPs (maxsnps), insertions (maxinss), insertion rate (insrate), deletions (maxdels), deletion rate (delrate), substitutions (maxsubs), and substitution rate (subrate). For “Perfect” and “Uneven” datasets, all values for controlling errors were set to 0 and an additional argument controlling inclusion of errors was set to false (adderrors=f). “True” datasets used the following values for error-controlling parameters: maxsnps=3, maxinss=2, maxdels=2, maxsubs=2, insrate=0.00000315, delrate=0.000005, subrate=0.0033, adderrors=t. The chosen error rates are based off average error rates observed by Schrimer and colleagues in their paper on Illumina™ error profiles from metagenomic data (Schirmer et al. 2016). Simulated reads for each genome of a metagenome were concatenated. Then, to induce random ordering of reads, the tool shuffle.sh from BBMap was used with default parameters. Read pairs were then separated into two FASTQ files using BBMap’s reformat.sh tool, specifying the single, shuffled, interleaved file be split into R1 and R2 FASTQ files. No filtering was performed on short reads as all reads possessed a q-score of 36.

### Metagenome assembly and quality assessment

Long read assembly was performed using Flye version 2.9-b1768 (Kolmogorov et al. 2020). Default parameters for Flye were used with the addition of ‘--nano-raw’ and ‘--meta’ to denote assembly of uncorrected reads from a metagenomic sample. SPAdes version 3.15.5 was used for metagenomic assembly of simulated short read data (Nurk et al., 2017). Default options for SPAdes were used except for the following: ‘--meta’ was specified to indicate the sequence data were from a metagenomic sample, kmer sizes (-k) consisted of a list of integers: 21, 33, 55, 77, 99, 127, ‘--phred-offset’ was set to 33 due to simulated short reads not possessing defined signatures of the PHRED quality, and ‘--only-assembler’ was specified to disable read error correction. Metagenomic assembly quality was evaluated using metaQUAST version 5.0.2 (Mikheenko, Saveliev, and Gurevich 2016). Default parameters were used except for supplying the directory containing the reference genomes for each metagenome to ‘-r’.

### Metagenome-assembled genome recovery and taxonomic classification

Metagenome-assembled genomes (MAGs) were generated using the following steps. First, long reads were mapped back to the assemblies using minimap2 version 2.24-r1122. For long reads, the flag ‘-x’ was given the ‘map-ont’ argument to optimize alignment parameters for nanopore sequence data (Li 2018). Short read headers were shortened using the tool seqtk-1.4 version r122’s ‘rename’ function (Li, 2023). Short reads were then mapped using bwa-mem2 version 2.2.1 by first creating an index from a metagenome’s assembly file using ‘bwa-mem2 index’, and then mapping the reads back using ‘bwa-mem2 mem’. Each alignment process produced an unsorted SAM file which was then converted into sorted BAM files using Samtools version 1.16.1 with options ‘samtools view -b’ for creating the BAM file, and ‘samtools sort’ for sorting the BAM file (Danecek et al. 2021). A coverage file was generated for each read type using CoverM version 0.6.1 using the default ‘-m mean’ for coverage estimation (Woodcroft 2021). For long reads, the mapper option for CoverM was set to ‘-p minimap2-ont’, and for short reads it was ‘-p bwa-mem’. Each coverage file was used to bin each read type’s respective assemblies using MetaBAT 2 version 2.15 with default parameters except for the addition of the ‘--cvExt’ argument to denote coverage files originated from a third-party tool (Kang et al. 2019). Binned contigs were then assessed for completeness and contamination using CheckM2 version 1.0.1 with default parameters (Chklovski et al. 2022). Taxonomic classification was performed using Kraken2 version 2.1.2 for both short and long read metagenomic assemblies (Wood, Lu, and Langmead 2019). Default parameters were used with the ‘--use-names’ flag included. The kraken2 report and output files were kept for further analysis.

### Result evaluation

For visual assessment of metagenome assembly quality, assembly graphs were visualized using Bandage version 0.8.1 (R. R. Wick et al. 2015). Resulting values for genome fraction recovery, NGA50, and number of misassemblies from metaQUAST were analyzed using the *statannotations* python package (Charlier et al., 2022). Statistical significance was evaluated using a two-sided Mann-Whitney U test with Bonferroni correction. A p-value threshold of 0.05 was used to determine significance. Performance evaluation of taxonomic classification from reads and assemblies was accomplished by calculating precision, recall, and F-score. Precision was defined as the rate of successful taxonomic assignment of a read/contig to the correct taxon within the metagenome over the total number of taxonomic assignments. Also referred to as the false positive rate, the 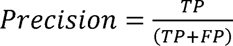 where TP is the true positive rate and FP is the false positive rate, or an incorrect classification of a read/contig at a given taxonomic rank. Recall, also defined as the rate of false negatives, was determined using the equation 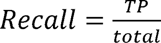 where ‘total’ is the expected number of distinct organisms at a specified taxonomic rank. F-score, or the harmonic mean, generates an overall score factoring in both precision and recall. F-score was calculated using the equation 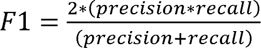 . A statistical comparison of precision, recall, and F-score was performed using a one-way ANOVA for assemblies and Mann-Whitney U test with Bonferroni correction for reads with the *statannotations* python package. A p-value threshold of 0.05 was used to determine significance.

Estimation of relative abundance of taxa at the genus and species level was carried out using Bracken version 2.8 (Lu et al., 2017). Short and long reads were processed through Kraken2 to generate report files that were then used as inputs for Bracken. Bracken then returns a report file containing estimates of relative abundance. For each genome in a metagenome, the predicted abundance was plotted against the actual abundance in a scatterplot along with a linear regression line using a linear regression function from SciPy version 1.10.1 (Virtanen et al., 2020). R-values, slope, and p-values for each read type’s line were reported.

MAG recovery rates were determined by comparing the total number of MAGs recovered by the binning of contigs from short and long read metagenomic assemblies. A MAG’s quality is determined by the completeness and contamination values of contigs assigned to a bin by MetaBAT2 and assessed by CheckM2. CheckM2 defines completeness as the percentage of universal, bacteria-specific gene markers present in a given bin. Contamination for CheckM2 is based on how many of these markers exist in multiple copies which indicates the presence of contigs that do not belong in a particular bin. For the minimum threshold of a bin to be considered a MAG, the cutoffs were defined as a completeness ≥50% and contamination ≤10% (A. Murat Eren 2016). For each of the 9 sets of metagenomes, the average number of MAGs recovered from short and long read assemblies was recorded. A Mann-Whitney U Test with Bonferroni correction was applied using the *statannotations* python package to compare differences in MAG recovery rates between short and long read assemblies. A p-value threshold of 0.05 was used to determine significance. Bins determined to be MAGs were also evaluated on quality which was defined by completeness only. High quality MAGs had a completeness ≥90%, medium quality MAGs had a completeness ≥70% and <90%, and low-quality MAGs had a completeness ≥50% and <70%, respectively.

### Comparison of experimental metagenomic data

To examine whether read type affects assessment of microbial composition, we generated short- and long-read sequencing data from 19 mouse fecal pellets collected during ongoing experimental studies. Long-read sequencing data were generated using the ONT GridION. Prior to DNA extraction, host DNA was depleted using an enzymatic approach as described previously with some modifications (Charalampous et al., 2019). Each sample extraction used 1-2 mouse fecal pellets. Fecal pellets were homogenized in 1 mL of InhibitEX buffer (Cat. No./ID: 19593, Qiagen, Hilden, Germany). After pelleting the homogenate and aspirating off the liquid, 250 µL of 1X PBS and 250 µL 4.4% saponin was added and mixed thoroughly before resting at room temperature for 10 minutes. Nuclease-free water (350 µL) along with 12 µL of 5M NaCl were added to induce osmotic lysis. After pelleting, the liquid was again aspirated off and the pellet was resuspended in 100 µL of 1X PBS. One-hundred microliters of HL-SAN enzyme buffer added (5.5_JM NaCl and 100_JmM MgCl2 in nuclease-free water), followed by 10 µL of HL-SAN endonuclease (Article No. 70910-202, ArcticZymes, Tromsø, Norway) before being incubated on a shaking incubator at 37°C for 30 minutes at 1000 rpm. Two final washes of the pellet were performed using 800 and 1000 µL of 1X PBS before proceeding to extraction. Genomic DNA (gDNA) was extracted and purified using the New England Biolabs Monarch® Genomic DNA Purification Kit (New England Biolabs, Ipswich, MA) following manufacturer instructions. Quality and concentration of gDNA was assessed using Agilent 4200 TapeStation System and Qubit 4 Fluorometer. Sequencing libraries for nanopore sequencing were prepared using the Rapid PCR Barcoding Kit (SQK-RPB004). Using an Oxford Nanopore GridION™, libraries were sequence on a R9.4.1 flow cell generating an average of 1.84 Gbps of sequence data with a mean read length of 3,491.62 bps and a mean read quality of 12.9. The longest read produced was 42,266 bps. Short read sequencing was performed by SeqCenter LLC (SeqCenter, Pittsburgh, PA) using an Illumina™ NovaSeq™ with Nextera Flex Library Preparation Kit® and V3 flow cell chemistry to produce an average of 2.53 Gbps of paired-end 2x150 bp reads across all samples. Illumina data were filtered using Trimmomatic v0.39 for paired-end reads with the following trimming settings: SLIDINGWINDOW:4:20 MINLEN:100.

Analysis followed the methodological approaches described previously for short- and long-reads, respectively. Reads were first classified using Kraken2 and relative abundances were estimated using Bracken. Before analysis, bracken reports were trimmed to remove all taxonomic identifications where the number of reads assigned to an organism fell below a given threshold of 1/20,000 (0.00005). To our knowledge, no studies using shotgun metagenomic sequencing have given the expected taxonomic diversity of the mouse gut microbiome at the genus or species level. The Murine Microbiome Database consists of 1732 genera and 4703 species, however these data come from 16S sequence analysis (Yang et al., 2019). An alternative database, the Comprehensive Mouse Microbiota Genome (CMMG), which derived 1,573 species from mouse gut metagenomes combined with known reference metagenome-assembled genomes at the time of its creation (Kieser et al., 2022). In this study, we empirically defined a cutoff where the count of unique taxa was closer to that of the CMMG, as 16S sequence often reports higher diversity. Initial plotting of the number unique taxa showed counts of over 1750 genera and over 6000 species for both short- and long-read data when no abundance cutoff was applied. Using a cutoff of 1/20,000 yielded unique taxa counts for species similar to that of the CMMG. Short- and long-read data for a group were then compared to each other using the beta_diversity.py script from the KrakenTools suite. Qualitative assessment was performed by generating a hierarchically clustered heatmap of the resulting Bray-Curtis Dissimilarity matrix. For quantitative analysis, a principal component analysis (PCA) was performed. First, data were transformed using a center-log ratio transformation (Aitchison, 1986). Then, PCA was run using the PCA function from scikit-learn’s decomposition library (Pedregosa et al., 2011). Briefly, sample IDs were separated as features and abundance values for each taxon at the genus and species level were retained. Abundance values for unique taxa were then given to the PCA object which created a fit and subsequently applied dimensional reduction to the abundance data. The resulting principal components were kept in a table from which the top two principal components were plotted.

## Results

### Comparison of metagenomic assembly completeness

Metagenomic assemblies for each synthetic dataset were qualitatively assessed through visualization (Figure S1). Long read assemblies produced graphs with greater contiguity and often resulted in circularization of unitigs, an indication they represented complete chromosome or, in the case of smaller, circularized contigs, plasmids.

Assemblies from long reads demonstrated significantly higher genome fraction coverage than their short read counterparts (Figure 2A, B, and C; Supplementary Table S1). This was observed across all metagenome sets, regardless of complexity, except for the synthetic dataset composed of 50 organisms with perfect reads (Figure 2A). Similar results were observed when comparing assemblies by NGA50 values, with long read data showing significantly higher scores (Figure 2D, E, and F; Supplementary Table S1). With “Perfect” data, short reads produced significantly fewer misassemblies, however this difference was not significant for larger “Uneven” and all “True” datasets (Figure 2G, H, and I; Supplementary Table S1).

**Figure 2.**
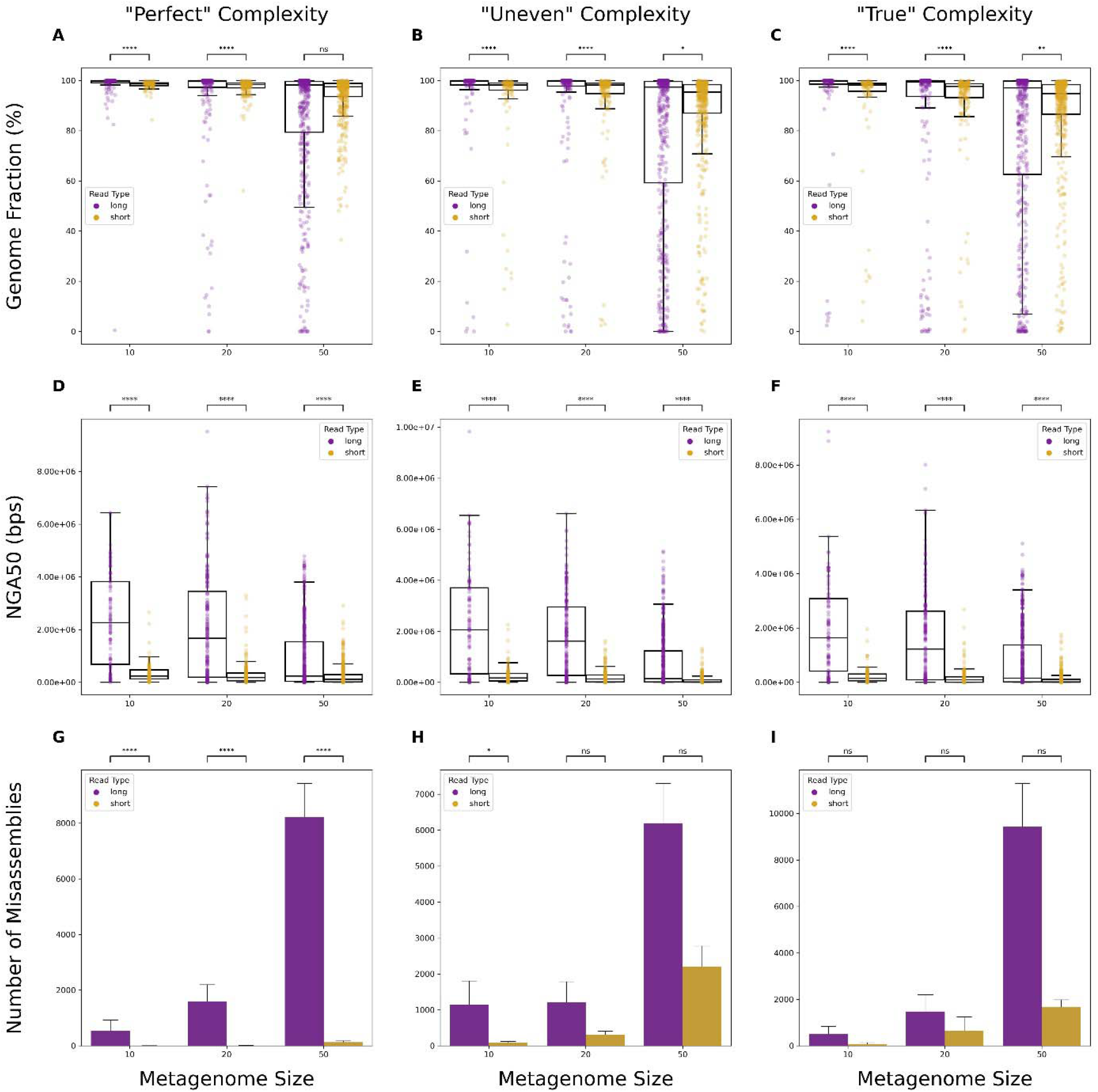
Comparisons of assembly quality metrics. Boxplots depicting differences in genome fraction values (A, B, C) and NGA50 (D, E, F) from all genomes in a set of metagenomes. Bar plots depicting total misassembly counts across all genomes in a set of metagenomes (G, H, I). “Perfect” denotes simulated data have no errors and abundances are evenly distributed for each organism in the metagenome. “Uneven” denotes simulated data have no errors, but variable abundances for each organism. “True” denotes simulated data have simulated sequencing errors based on the read type, as well as variable abundances for each organism. Significance was calculated using a Mann-Whitney U Test with Bonferroni correction. ns: non-significant p-value (p > 0.05), *: p ≤ 0.05, **: p ≤ 0.01, ***: p ≤ 0.001, ****: p ≤ 0.0001.

### Evaluating taxonomic classification

Evaluation of taxonomic classifications using assemblies at the genus and species-level was assessed for precision, recall, and F-score. Neither read types significantly differed when measuring recall of taxa (Figure 3D, E, and F; Figure 4D, E, and F). However, long reads significantly outperformed short reads in precision across almost all levels of metagenome complexity at both the genus and species level (Figure 3A, B, and C; Figure 4A, B, and C). The F-score, or harmonic mean of the precision and recall, was calculated to measure the accuracy of classification results using either short or long reads. Long reads demonstrated significantly higher F-scores than short reads across almost all levels of complexity, indicating that long reads provide greater accuracy for taxonomic classification.

**Figure 3.**
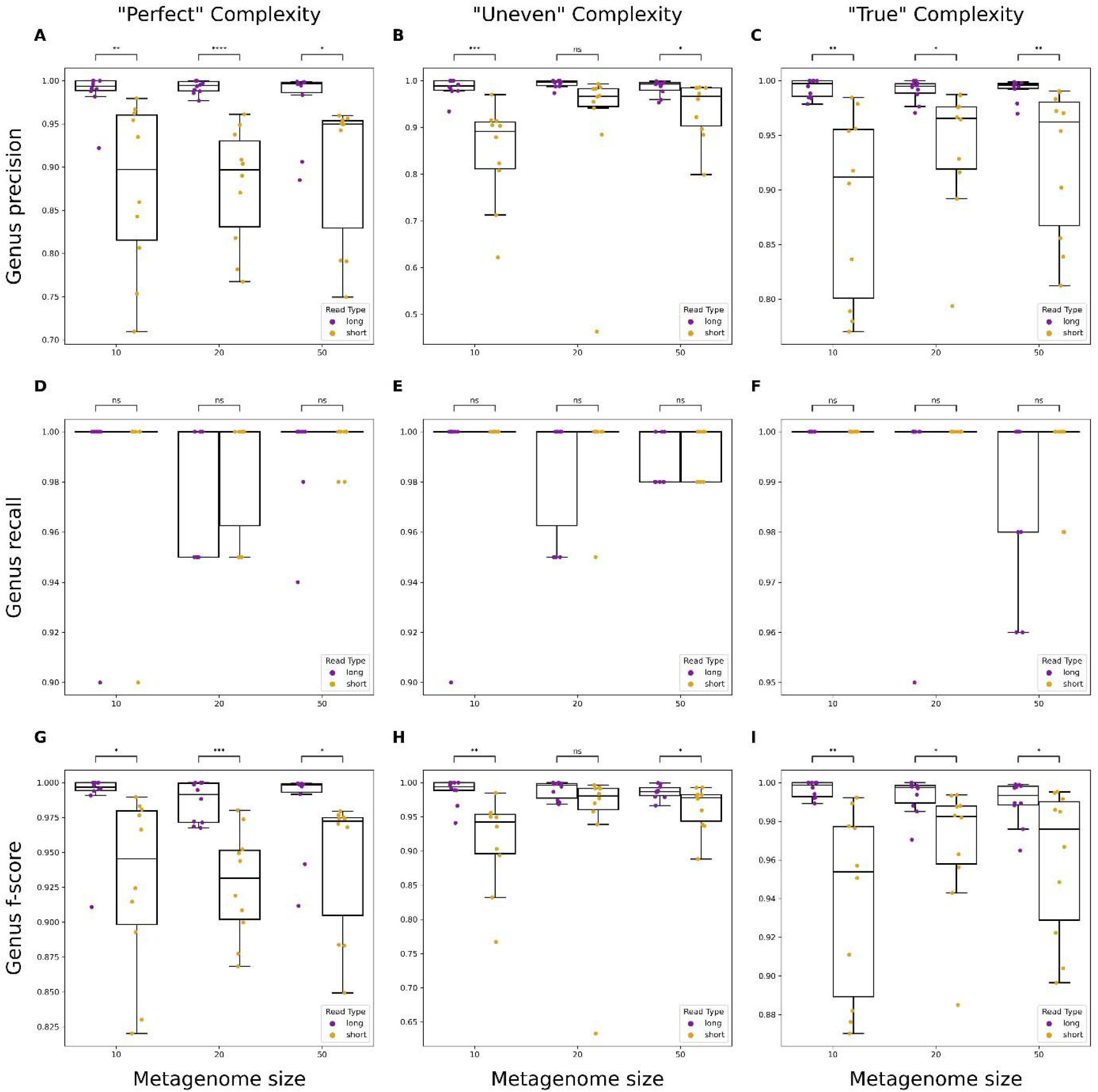
Performance evaluation of genus-level classification on metagenomic assemblies using short or long reads. Boxplots showing precision, recall, and F-score metrics for metagenomes of varying size and complexity. “Perfect” simulated reads have no errors and even abundances across organisms (A, D, G), “Uneven” have no errors and randomly varied abundances across organisms (B, E, H), and “True” have simulated errors specific to each read type and varied abundances across organisms (C, F, I). Significance was calculated using a One-way ANOVA. ns: non-significant p-value (p > 0.05; blank means p = 1), *: p ≤ 0.05, **: p ≤ 0.01, ***: p ≤ 0.001, ****: p ≤ 0.0001.

**Figure 4.**
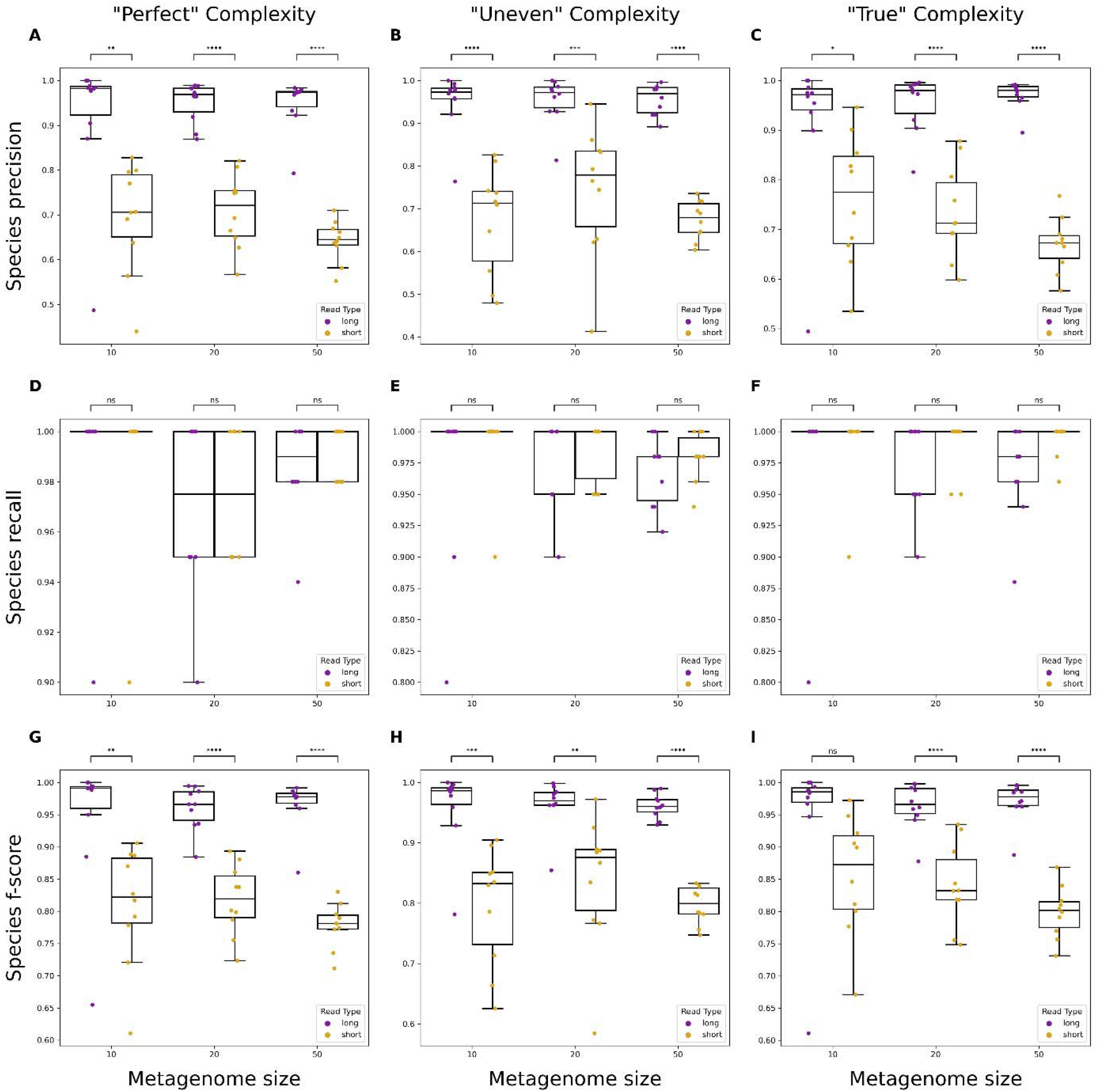
Performance evaluation of species-level classification on metagenomic assemblies using short or long reads. Boxplots showing precision, recall, and F-score metrics for metagenomes of varying size and complexity. “Perfect” simulated reads have no errors and even abundances across organisms (A, D, G), “Uneven” have no errors and randomly varied abundances across organisms (B, E, H), and “True” have simulated errors specific to each read type and varied abundances across organisms (C, F, I). Significance was calculated using a One-way ANOVA. ns: non-significant p-value (p > 0.05; blank means p = 1), *: p ≤ 0.05, **: p ≤ 0.01, ***: p ≤ 0.001, ****: p ≤ 0.0001.

### Estimating relative abundance using short and long read data

Before analyzing the performance of each read type when used for relative abundance estimation, the classification capabilities of the reads were first assessed. Precision, recall, and F-score of genus-level classification was not significantly affected by read type in most cases, which differed from classification using metagenomic assemblies (Supplementary Figure S2). However, the performance for each read type on species-level classification was significantly improved when using long reads (Supplementary Figure S3). Both precision and F-score were higher for classifications using long reads and recall was not significantly different between the two. Examining accuracy in estimation of relative abundance found long reads outperformed short reads in most cases at the genus and species level (Figure S4; Figure 5).

**Figure 5.**
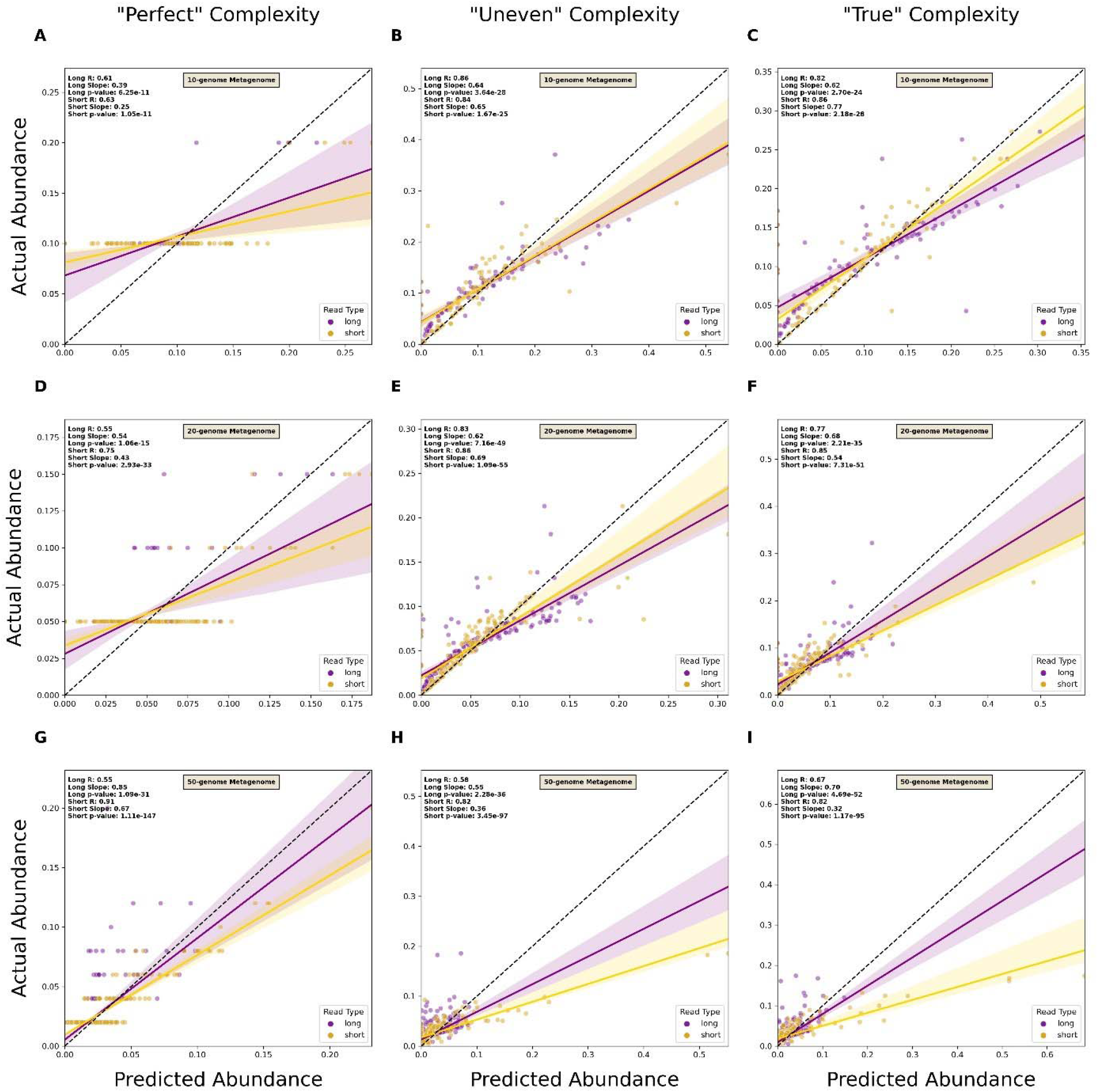
Comparison of short and long read capacity for species-level relative abundance estimation. Scatterplots of predicted versus actual abundance values for genomes present in simulated metagenomes from short and long read data. Read types were compared across metagenomes of varying complexity at both the genus and species level. “Perfect” datasets consisted of reads without sequencing errors and organism abundance was evenly distributed (A, D, G), “Uneven” consisted of reads without sequencing errors and randomly distributed abundances of organisms (B, E, H), and “True” consisted of reads with simulated errors and randomly distributed abundances of organisms (C, F, I). A linear regression line was plotted for each read type. Each line has its reported R-value, slope, and p-value. The dotted line represents the 1-to-1 line where values of predicted abundance match values of actual abundance.

### Metagenome-assembled genome (MAG) recovery from short- and long-read data

In most cases, neither read type resulted in more MAGs being recovered except for short reads producing more MAGs from “True” 50-genome metagenomes (Figure 6C). It was found that the binning of contigs from long read assemblies generally produced more total bins than assemblies produced from short reads (Supplementary Table S2). Following this observation, the quality of MAGs recovered were evaluated. Bins were assigned a quality of high, medium, or low, depending on their completeness. Long reads did not result in more high-quality MAGs at any level of complexity; however, they did show significantly more medium and low-quality MAGs in some cases (Supplementary Figure 5D, G, H). Complementing Figure 6, short reads recovered significantly high-quality MAGs from the most complex datasets (Supplementary Figure 5I).

**Figure 6.**
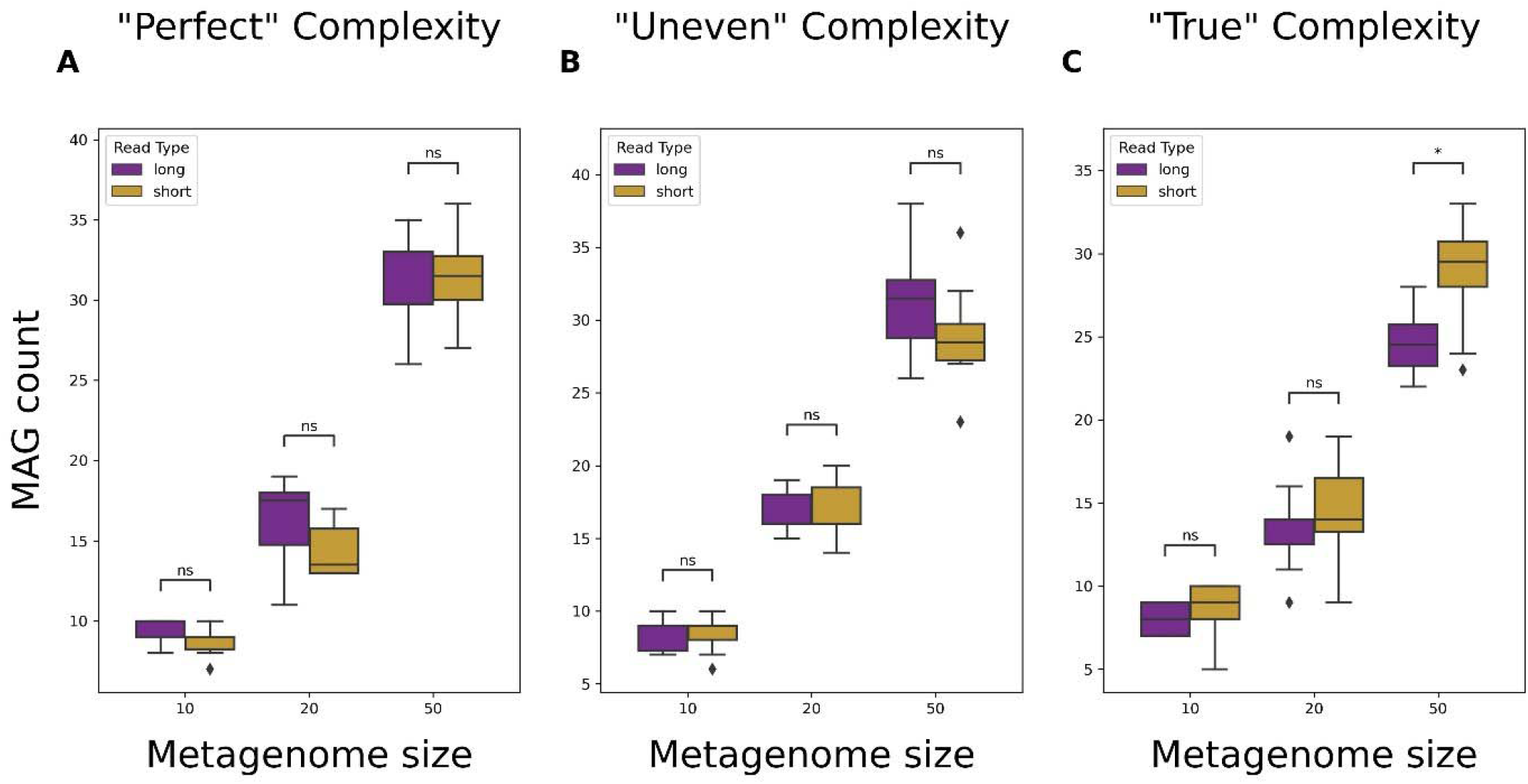
Comparison of metagenome-assembled genome (MAG) recovery capabilities from assemblies produced using short or long reads. Total number of MAGs recovered from assemblies of simulated short or long read sequence data. “Perfect” datasets consisted of reads without sequencing errors and organism abundance being evenly distributed (6A), “Uneven” consisted of reads without sequencing errors and organism abundance being randomly assigned (6B), and “True” consisted of reads with simulated sequencing errors and organism abundance being randomly assigned (6C). Significance was calculated using a Mann-Whitney U Test with Bonferroni correction. ns: non-significant p-value (p > 0.05), *: p ≤ 0.05, **: p ≤ 0.01, ***: p ≤ 0.001, ****: p ≤ 0.0001.

### Microbial compositional differences between short and long reads

Using experimental data obtained from mouse fecal pellets, we examined whether read type significantly altered inferences of microbial composition for a metagenomic sample. Qualitative assessment showed clustering of results around similar read types instead of around sample types (Figure S6). This trend was also observed when results were run through a PCA where samples clustered around similar read types instead of around similar sample types (Figure 7).

**Figure 7.**
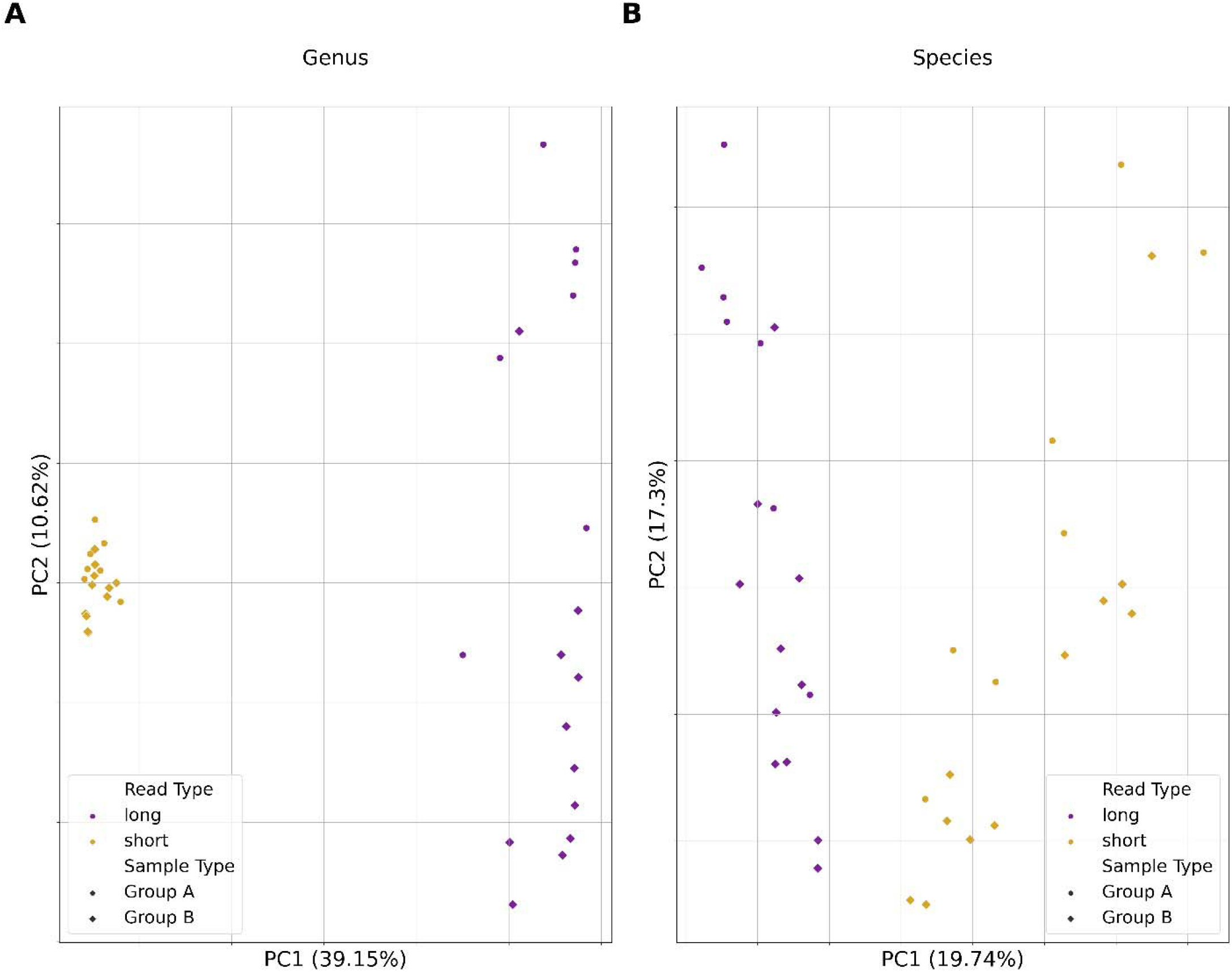
Principal Component Analysis (PCA) results from short and long read sequence data analysis of empirical metagenomic samples from mouse fecal pellets. PCA plot depicting the distribution of metagenomic samples according to their two main principal components. Coloration indicates the read type: purple for long read data and yellow for short read data. Shape indicates the sample type. Principal components 1 and 2 are on the x and y-axis respectively with their explained variance ratios written as a percentage.

## Discussion

For metagenomic studies of microbial communities, long-read sequencing platforms potentially offer numerous advantages, yet short read sequencing remains the most commonly applied technology (Watson et al. 2021). We sought to quantitatively assess the use of long-read sequence data by using simulated metagenomes with increasing complexity. Evaluation of read type performance across several metrics showed that long and short reads possess respective strengths. We also used paired long and short read metagenomic data from murine fecal samples to assess the generalizability of our findings. Overall, we highlight that choice of sequence type has a direct impact on the ability to accurately assemble metagenomes, identify comprising members of a diverse microbial community, and recover MAGs from metagenomic sequence data.

For optimal performance, sequencing data must be capable of producing high quality assemblies for taxonomic classification, relative abundance estimation, functional annotation, and other related analyses. We show that in simulated long read assemblies had significantly higher percent genome coverages than short read assemblies. Analyses that rely on examining contiguous sequences, such as gene annotation, would thus benefit from assemblies generated using long-read data. Short reads are known to struggle with repetitive regions that are common in many bacterial genomes (Hu et al., 2021), which likely explains the lower genome coverages and greater discontiguity observed from short read assemblies. Our results also suggest long reads would perform better at recovering novel genomes; however, that was not directly tested here. Higher NGA50 values indicate greater contiguity, further supporting the findings highlighted by higher genome fraction observed. Regarding misassembly rates, short reads did produce fewer misassemblies in all “Perfect” datasets (Figure 2G). However, in more complex datasets, the observed differences were found to not be significant in most cases (Figures 2H, I). One explanation is that metagenomic assemblers attempt to balance accuracy and contiguity, thus increasing the possibility of misassemblies (Kolmogorov et al., 2020). The lower per-base accuracy of long-read sequencers has long been a reason researchers tend to favor short reads. Our results highlight the shrinking gap in result quality of assemblies produced with either short or long reads.

Taxonomic classification is a central to metagenomic studies and several tools exist for use on filtered reads or assembled contigs. Previous studies have compared classification using partial and full 16S sequencing using short and long read platforms respectively (Hahn et al., 2016). Here, we demonstrate the capabilities of each read type when classifying whole metagenomic sequence data. For long-read assemblies, we found that classifications demonstrated significantly higher precision at both the genus and species level overall. The greater precision likely stems from long read assemblies being more contiguous, thus having fewer small fragments that could be improperly classified like in short read assemblies. When classifying reads at the genus level, both long and short reads performed similarly (Figure S2). This was partly explained by long reads having a lower precision compared to classifications with contigs. This is expected given there are more reads than contigs, thus more chances for reads to be misassigned to an improper taxon. When analyzing reads at a finer taxonomic resolution, long reads performed significantly better. For differentiation of species within a genus, larger reads provide more information for distinguishing the genome of origin. For assemblies and filtered reads, we observed no difference in sensitivity. These findings are relevant for the potential application of long-read metagenomic sequencing to infectious disease diagnostics (Bastida et al., 2021; Hoang et al., 2022; Huang et al., 2019). For example, the ability of ONT sequencing platforms to generate sequencing reads in real time, which can be used for taxonomic classification without sacrificing precision or sensitivity, is invaluable and avoids lengthy culturing, isolation, and molecular diagnostics.

In addition to taxonomic classification, assessment of differences in the relative abundance of comprising species is of great interest. This generally involves alignment of reads to a reference genome database and subsequent estimation of abundances from the coverage, which can be performed using both short and long reads (Verma & Sharma, 2020; Warwick-Dugdale et al., 2019). Our results showed that predicted abundances from long read data were more likely to be close to the actual abundance data for both genus and species-level classification (Figure 5; S4). This was likely due to the role of Kraken2’s results in Bracken’s estimation of organismal relative abundances. Bracken takes the initial classifications and then attempts to re-assign reads to proper taxa using a Bayesian-based approach. Having greater accuracy in these classifications enables Bracken to more likely assign reads to taxa accurately. Assessment of relative abundance thus benefitted from the higher precision of long reads. Research that derives conclusions based on significant shifts in abundances of all taxa or specific organisms can thus benefit from reliance on abundance estimation. Also, in clinical diagnostics, accurate estimation of abundance may enable clinicians to identify the etiologic microbe associated with the disease of interest (Wen et al., 2017).

A considerable challenge in microbial research is that most bacteria are non-culturable, which makes identification of organisms of potential importance difficult. Using shotgun sequencing and *de novo* metagenomic assembly, it is possible to recover both known and novel genomes and infer the identity and functional capacities of the organisms (Parks et al. 2017). Assemblies produced from either read type resulted in similar MAG recovery rates except short reads did identify more MAGs in “True” 50-genome metagenomes (Figure 6). A key tool in the binning process was MetaBAT2, which is highly dependent on assembly quality for performance. As a result, while long reads were more contiguous and often more complete, misassembly rates may have impacted read alignment during the MAG identification process. This is supported by our observation that long read assemblies contained more misassembles, even though misassembly rates were not significantly different between read types (Supplementary Table S2). This likely resulted in more bins with fewer contigs spread among them and therefore lower completeness due to the cutoffs used for MAG reporting. This also explains why long reads produced at times more medium or low-quality MAGs, since MAG quality was primarily determined by completeness, thus having contigs inadequately binned would result in lower completeness of MAGs.

Selection of sequencing platform has previously been shown to directly impact the taxonomic assignment of partial/total 16S rRNA sequences (Stevens et al., 2023). In the present study, bacterial metagenomic DNA extracted from murine fecal samples were sequenced using both short- and long-read sequencing platforms. Qualitative assessment measuring the beta diversity among samples of the same type, but different sequence data, revealed samples clustered by sequence type, not sample type (Supplementary Figure 6). Comparisons using PCA revealed the same trend of samples clustering by sequence type, rather than by sample type (Figure 7). This is likely due to an inflation of misclassified taxa in short read assemblies, causing clustering to be associated with those unique taxa. This is consistent with our findings that simulated short read data produced more false positives than long reads. This, in turn, artificially increased the diversity of the sample and explains clustering due to read type instead of sample type.

Taken together, our results show that long read data offers certain advantages towards accurate representation of a metagenome. Specifically, long reads resulted in greater accuracy when taxonomically classifying contigs and reads, and their assemblies were more contiguous and returned greater genome fractions for genomes within a metagenome. While this study provides evidence supporting the use of long read sequencing for metagenomic research, it does have some limitations. The most complex metagenomes that were simulated consisted of 50 organisms with variable abundance and simulated sequencing errors specific to each read type, but empirical data may consist of hundreds or even thousands of unique organisms (Bastida et al. 2021). Future work could employ simulations that mimic high diversity as well as abundance distributions observed in real metagenomic samples; however, we feel that increasing taxa would only increase the disparity between sequence data types. Another consideration is that pipelines for processing and analyzing metagenomic data can vary considerably. Various tools exist for read correction, assembly, polishing, taxonomic classification, and binning, all of which can affect results. Steps such as read error correction and assembly polishing are often used in metagenomic studies; however, we did not include them here to reduce computational requirement and to limit variables in the comparison, especially if they are unique to one read type. Another limitation was that binning algorithms are usually designed with short reads in mind, leveraging the higher coverage of data that accompanies short read sequencing. In recent years, tools such as a LRBinner have emerged which offer better binning for error-prone long reads (Wickramarachchi and Lin 2022), but those were not applied here. Overall, we aimed to use the most commonly employed tools used for these studies at the time of writing. Last, we used a limited number of real-world samples, which could be expanded in future studies.

Short and long read sequencing have their respective places in metagenomic research. Short read sequencing does have its strengths, especially with pipelines for analyzing short read data being more established. Our results highlight that both read types perform comparably well across several metagenomic analyses, in particular assembly quality and MAG recovery. For studies concerned with misassemblies, such as those focused on specific species of interest, short reads produce significantly fewer and may be preferable. For studies concerned with microbial composition, long reads provided higher precision with similar sensitivity to short reads despite lower sequencing depths. They also provide better estimations of organismal abundance within metagenomic samples. Leveraging the strengths of both read types is also possible. Tools such as OPERA-MS combine long reads containing more genetic information per-read with high accuracy short reads (Bertrand et al., 2019). Other programs like PolyPolish enable the use of short reads for improving long read assembly quality (Wick & Holt, 2022). Further advancement may shift practice towards use of hybrid approaches as has occurred with genome sequencing. In conclusion, our results show long read sequence data outperforms short read sequence data when used for taxonomic classification and metagenomic assembly contiguity. This work provides evidence for the consideration of long read sequencing for research focused on accurate reconstruction of microbial populations and its strength in clinical diagnostics.

## Figure Legends

**Supplemental Figure 1.**
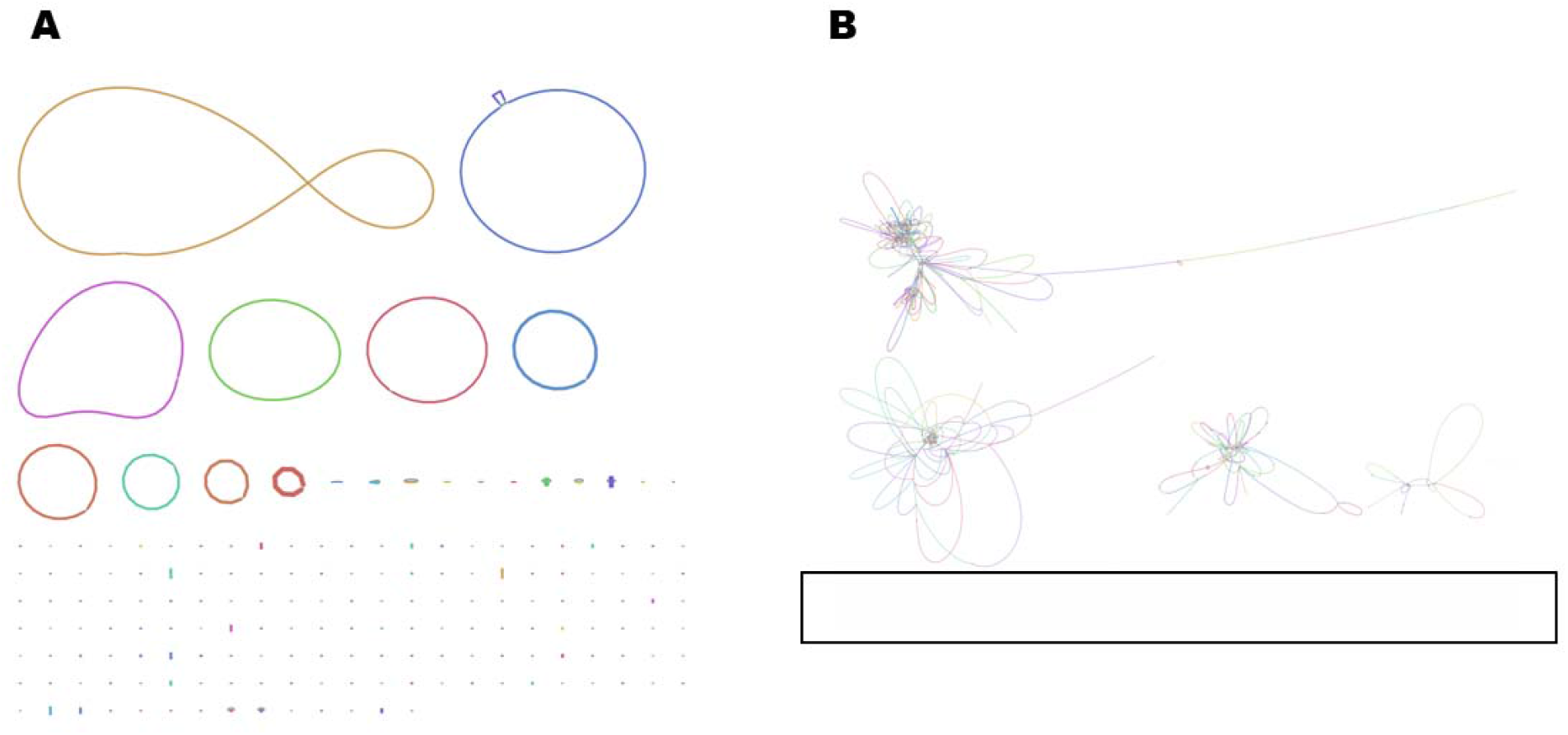
Assembly graphs from a “True” 10-genome metagenome visualized by Bandage. 2A. Bandage graph generated from metagenomic assembly of long read sequence data. 2B. Bandage graph generated from metagenomic assembly of short read sequence data. The black box highlights small, disconnected contigs.

**Supplemental Figure 2.**
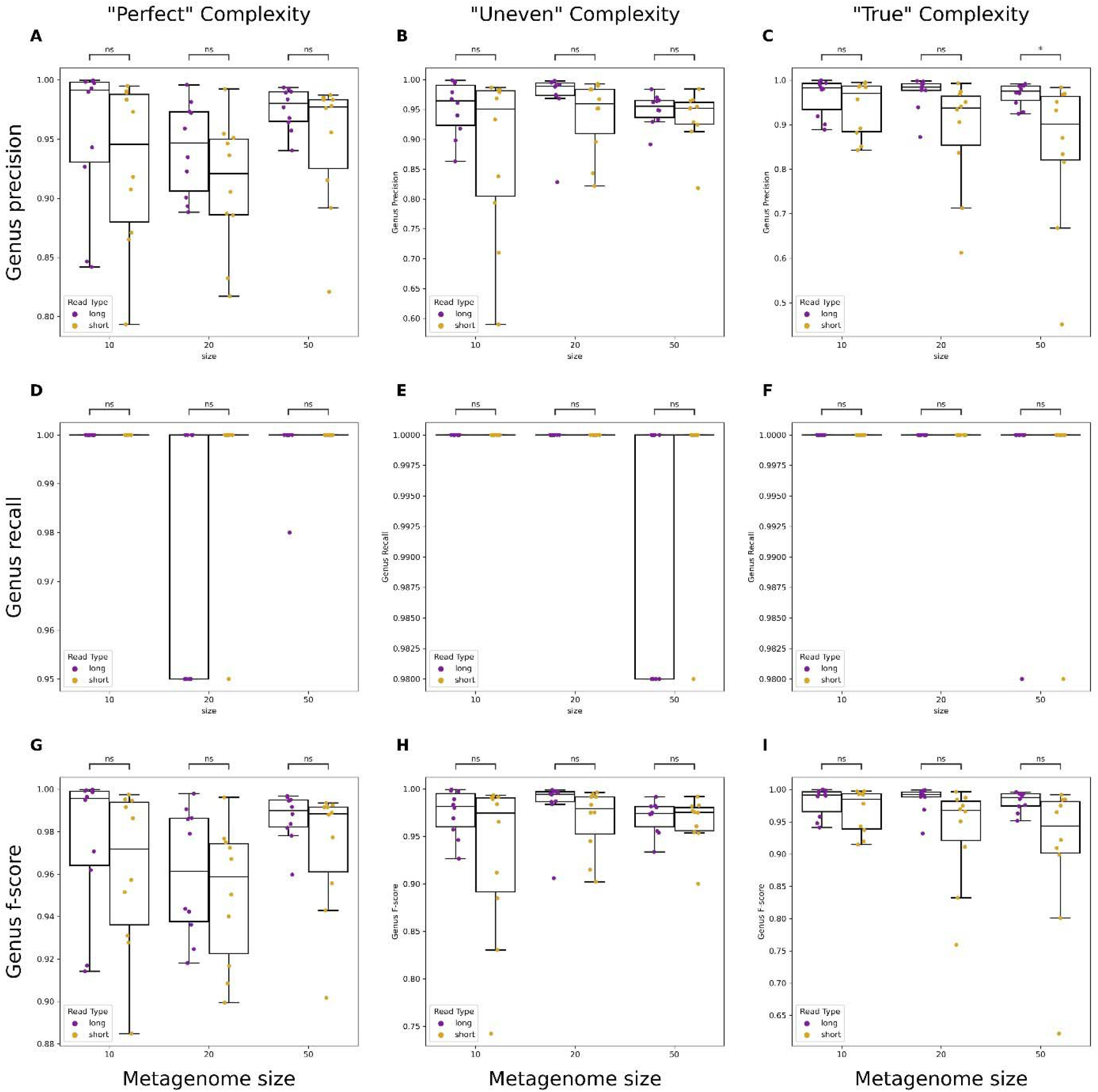
Performance evaluation of genus-level classification using short and long reads. Boxplots showing precision, recall, and F-score metrics for metagenomes of varying complexity. “Perfect” simulated reads have no errors and even abundances across organisms (A, D, G), “Uneven” have no errors and randomly varied abundances across organisms (B, E, H), and “True” have simulated errors specific to each read type and varied abundances across organisms (C, F, I). Significance was calculated using a Mann-Whitney U test with Bonferroni correction. ns: non-significant p-value (p > 0.05; blank means p = 1), *: p ≤ 0.05, **: p ≤ 0.01, ***: p ≤ 0.001, ****: p ≤ 0.0001.

**Supplemental Figure 3.**
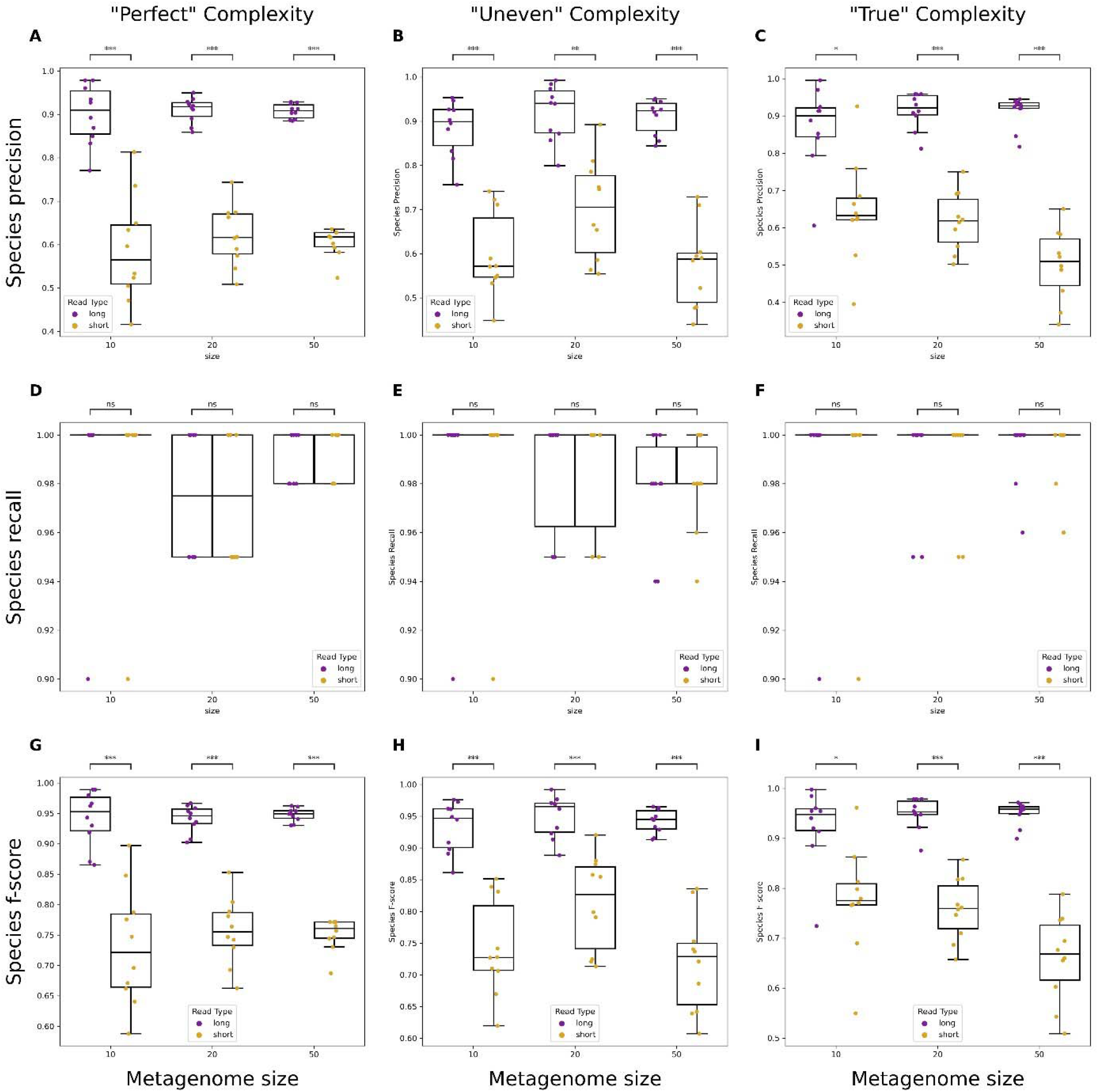
Performance evaluation of species-level classification using short and long reads. Boxplots showing precision, recall, and F-score metrics for metagenomes of varying complexity. “Perfect” simulated reads have no errors and even abundances across organisms (A, D, G), “Uneven” have no errors and randomly varied abundances across organisms (B, E, H), and “True” have simulated errors specific to each read type and varied abundances across organisms (C, F, I). Significance was calculated using a Mann-Whitney U test with Bonferroni correction. ns: non-significant p-value (p > 0.05; blank means p = 1), *: p ≤ 0.05, **: p ≤ 0.01, ***: p ≤ 0.001, ****: p ≤ 0.0001.

**Supplemental Figure S4.**
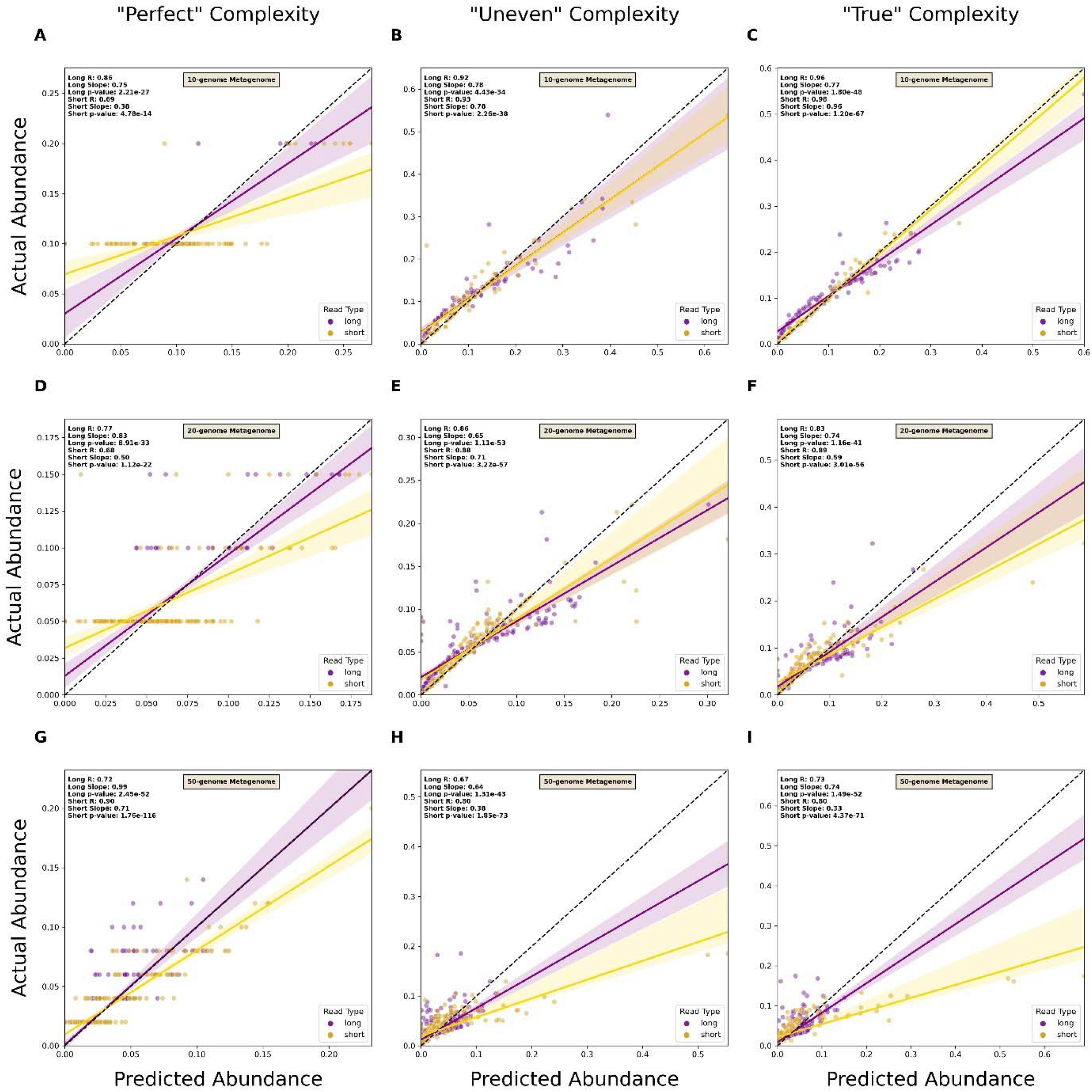
Comparison of short and long read capacity for genus-level relative abundance estimation. Scatterplots of predicted versus actual abundance values for genomes present in simulated metagenomes from short and long read data. Read types were compared across metagenomes of varying complexity at both the genus and species level. “Perfect” datasets consisted of reads without sequencing errors and organism abundance was evenly distributed (A, D, G), “Uneven” consisted of reads without sequencing errors and randomly distributed abundances of organisms (B, E, H), and “True” consisted of reads with simulated errors and randomly distributed abundances of organisms (C, F, I). A linear regression line was plotted for each read type. Each line has its reported R-value, slope, and p-value. The dotted line represents the 1-to-1 line where values of predicted abundance match values of actual abundance.

**Supplemental Figure 5.**
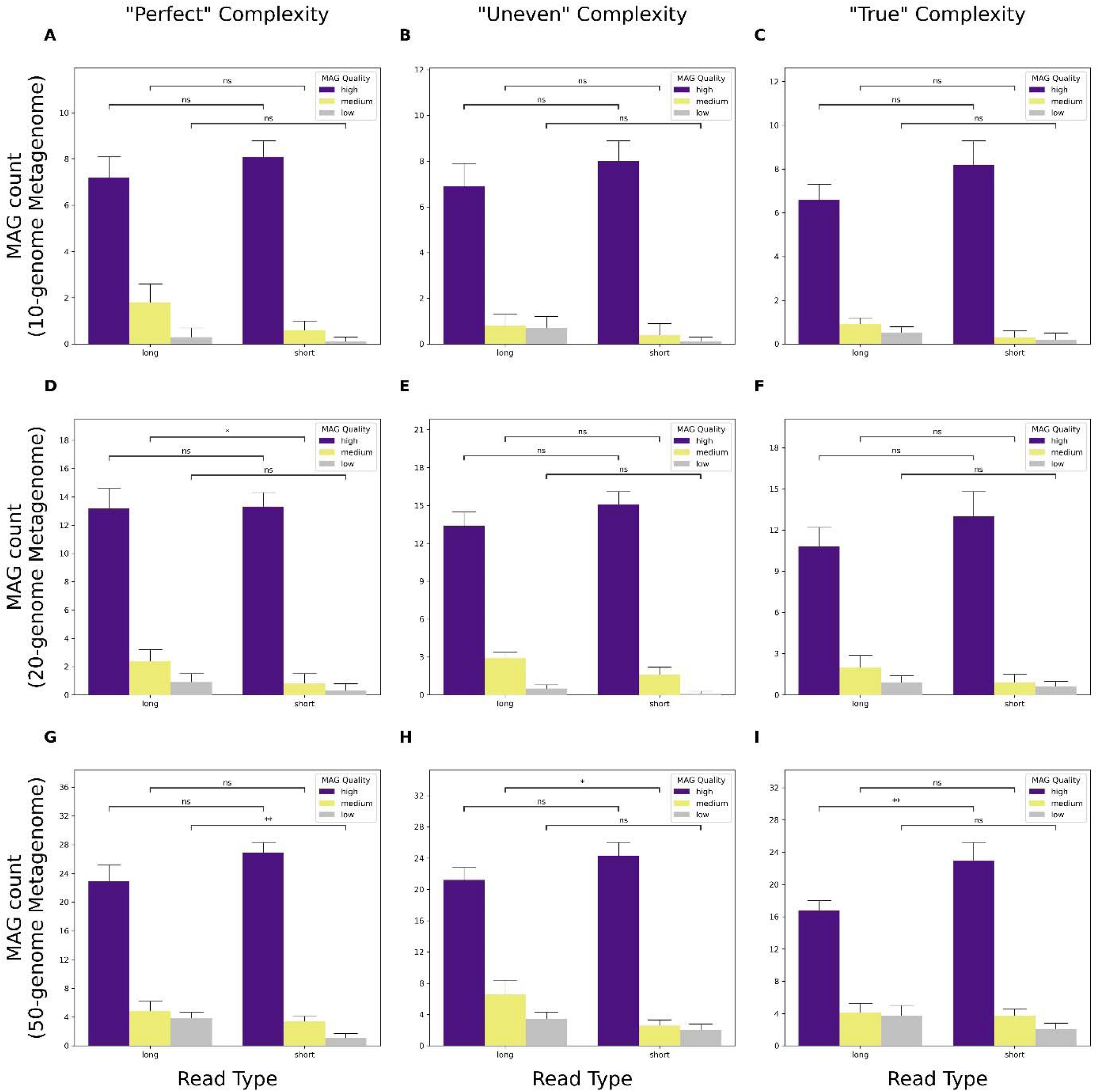
Comparison of metagenome-assembled genome (MAG) counts recovered from short and long read assemblies by quality. Barplots depict the total number of MAGs of a given quality recovered from both short and long read assemblies. “Perfect” denotes simulated data have no errors and abundances are evenly distributed for each organism in the metagenome (S4A-C), “Uneven” have no errors, but variable abundances for each organism (S4D-F), and “True” have simulated sequencing errors based on the read type, as well as variable abundances for each organism (S4G-I). Significance was calculated using a Mann-Whitney U Test with Bonferroni correction. ns: non-significant p-value (p > 0.05), *: p ≤ 0.05, **: p ≤ 0.01, ***: p ≤ 0.001, ****: p ≤ 0.0001.

**Supplemental Figure 6.**
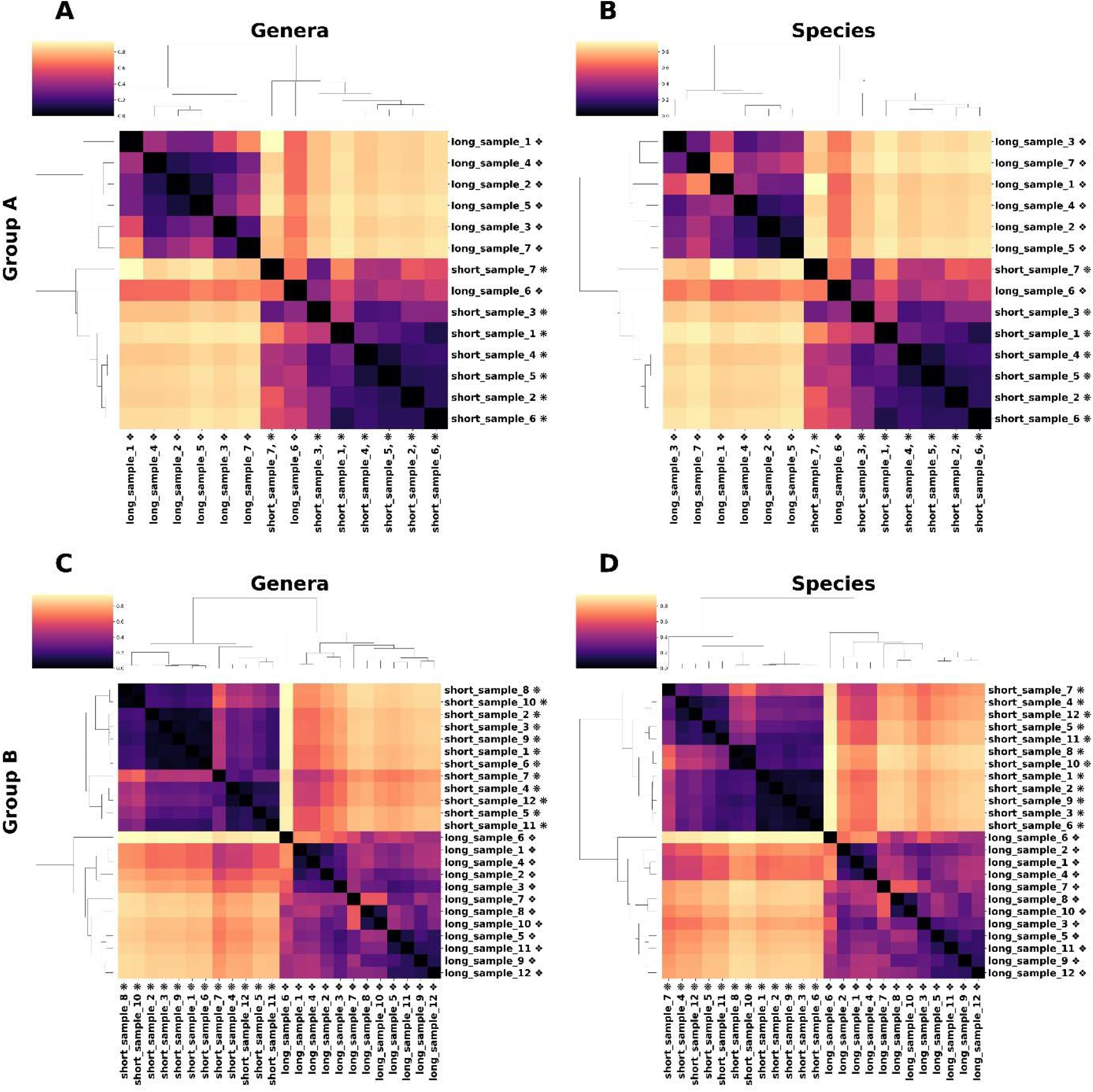
Hierarchically clustered heatmap of Bray-Curtis Dissimilarity. Heatmap visualizing the dissimilarity of samples based on the read type used in beta-diversity assessment. Brighter values indicate lower similarity (or higher Bray-Curtis dissimilarity), darker values indicate higher similarity (or lower Bray-Curtis dissimilarity). IZ: Samples consist of long read data. IZ: Samples consist of short read data

**Table S1.** Results of Mann-Whitney U testing for genome fraction, NGA50, and number of misassemblies. ^1^ “Perfect” indicates reads had no errors and abundances were evenly distributed. “Uneven” denotes reads had no errors, but variable abundances for each organism. “True” reads had simulated sequencing errors based on the read type and variable abundances for the organisms. ^2^P-value from Mann-Whitney U test with Bonferroni correction

| **simulated metagenome** | **complexity<sup>1</sup>** | **metaQUAST metric** | **statistic** | **p-value<sup>2</sup>** |
| --- | --- | --- | --- | --- |
| 10-genome metagenome | perfect | genome fraction | 7600.5 | 1.79E-10 |
| 10-genome metagenome | perfect | NGA50 | 8465 | 2.56E-17 |
| 10-genome metagenome | perfect | number of misassemblies | 6473 | 5.46E-08 |
| 20-genome metagenome | perfect | genome fraction | 27671.5 | 3.04E-11 |
| 20-genome metagenome | perfect | NGA50 | 30657.5 | 3.03E-20 |
| 20-genome metagenome | perfect | number of misassemblies | 29583.5 | 1.24E-26 |
| 50-genome metagenome | perfect | genome fraction | 134023.5 | 0.04813137 |
| 50-genome metagenome | perfect | NGA50 | 156277 | 7.40E-12 |
| 50-genome metagenome | perfect | number of misassemblies | 194570 | 3.23E-67 |
| 10-genome metagenome | uneven | genome fraction | 7433 | 2.59E-09 |
| 10-genome metagenome | uneven | NGA50 | 8152 | 1.35E-14 |
| 10-genome metagenome | uneven | number of misassemblies | 6008 | 0.0044121 |
| 20-genome metagenome | uneven | genome fraction | 28562.5 | 1.17E-13 |
| 20-genome metagenome | uneven | NGA50 | 32254 | 2.99E-26 |
| 20-genome metagenome | uneven | number of misassemblies | 20066 | 0.94863897 |
| 50-genome metagenome | uneven | genome fraction | 137225 | 0.00742218 |
| 50-genome metagenome | uneven | NGA50 | 169597 | 1.32E-22 |
| 50-genome metagenome | uneven | number of misassemblies | 133230.5 | 0.0645971 |
| 10-genome metagenome | true | genome fraction | 7812 | 5.66E-12 |
| 10-genome metagenome | true | NGA50 | 8341 | 3.26E-16 |
| 10-genome metagenome | true | number of misassemblies | 5768.5 | 0.01977121 |
| 20-genome metagenome | true | genome fraction | 26882.5 | 2.51E-09 |
| 20-genome metagenome | true | NGA50 | 29641.5 | 7.20E-17 |
| 20-genome metagenome | true | number of misassemblies | 20560 | 0.583517 |
| 50-genome metagenome | true | genome fraction | 138987.5 | 0.00218585 |
| 50-genome metagenome | true | NGA50 | 170176 | 3.94E-23 |
| 50-genome metagenome | true | number of misassemblies | 122815 | 0.62025684 |

**Table S2.**
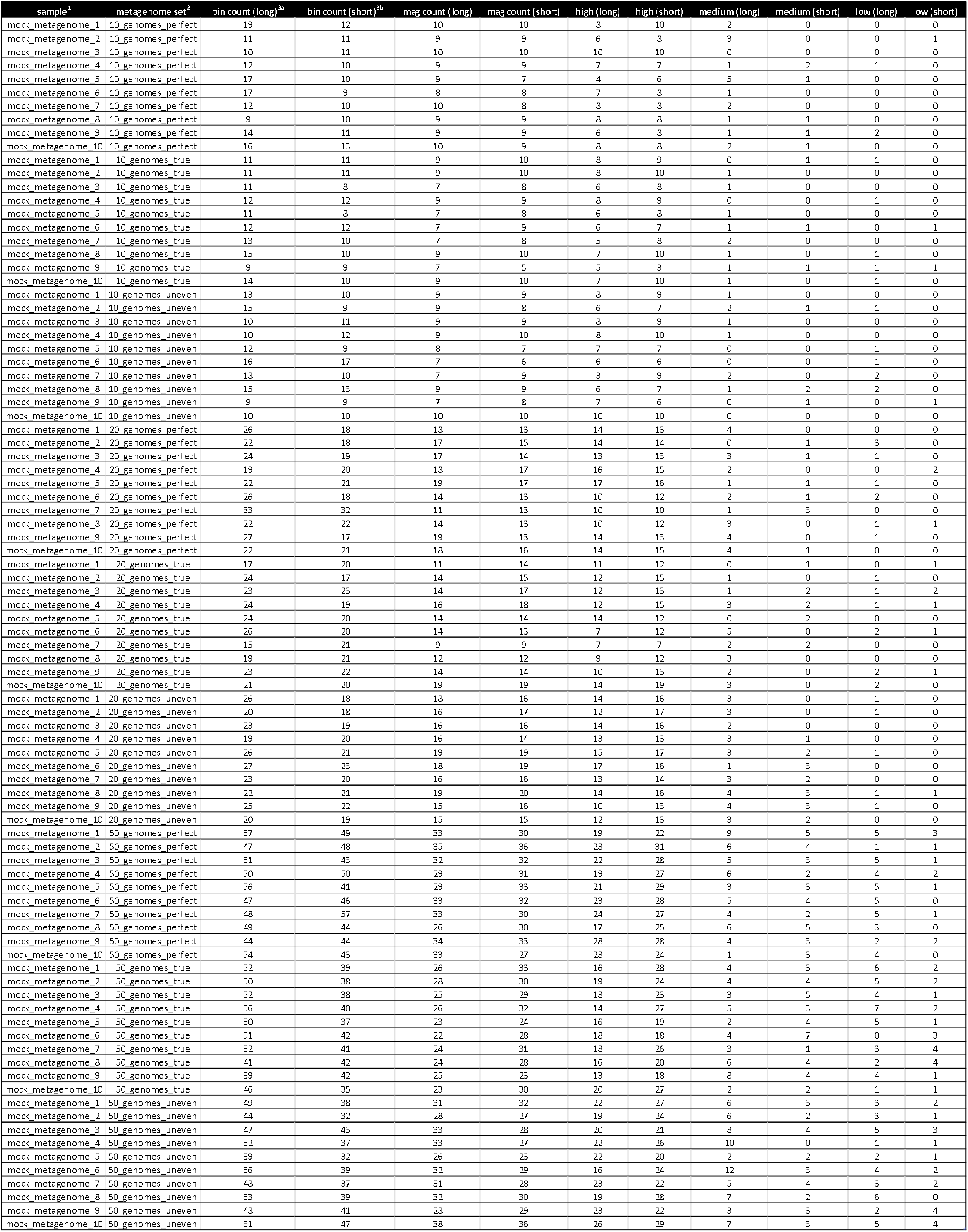
Bin and metagenome-assembled genome (MAG) counts from metagenomic samples. Total bins recovered for each metagenomic sample and counts of MAGs identified from those bins. MAGs were further separated by quality as described in the methods section. ^1^Samples consist of the 10 metagenomes simulated for a given complexity. ^2^A metagenome set is the given complexity level for a collection of simulated metagenomes. ^3a^Sum of all bins predicted by MetaBAT2 from long read assemblies. ^3b^Sum of all bins predicted by MetaBAT2 from short-read assemblies.

